# EGFR Mutation Subtypes Modulate Distinct Metabolic Profiles and Clinical Outcomes in Lung Cancer: A Retrospective Analysis

**DOI:** 10.1101/2025.06.26.659534

**Authors:** Tenzin Kungyal, Migmar Tsamchoe

## Abstract

Lung cancer continues to be a leading contributor of cancer-related mortality worldwide, with non-small cell lung cancer (NSCLC) as most prevalent cases. Identification of epidermal growth factor receptor (EGFR) mutations has profoundly enhanced our understanding and treatment of NSCLC, leading to the development of precision therapies, including EGFR tyrosine kinase inhibitors (TKIs). This retrospective study analyzed EGFR mutation distributions and their effect on overall survival (OS) using data accessed from TCGA. Our analysis revealed that EGFR mutations are most prevalent in lung cancer, with L858R appearing as the most frequent mutation, followed by E746_A750del and T790M. Remarkably, OS analysis exhibited that C797S mutations were linked with the least OS, with T790M, G719S, L861Q, and G719A also displaying significantly decreased OS compared to L858R mutations. Gene set enrichment analysis (GSEA) of T790M versus L858R cases revealed significant metabolic reprogramming in T790M mutants, noticeable by upregulation of oxidative phosphorylation (CPT1A, NDUFS1), lipid metabolism (HMGCR), and mTORC1 signaling. Metabolic adaptation observed in T790M indicates elevated bioenergetic flexibility and detoxification efficiency, possibly leading to therapeutic resistance. The work underscores EGFR mutation subtypes as distinct biological entities with distinctive metabolic needs, suggesting HMGCR (statin-targetable) and PPARα agonists as probable therapeutic paths for T790M-driven resistance. These understandings support for mutation-specific treatment approach and highlight the necessity to combine metabolic pathway targeting with EGFR blockade to augment responses in lung cancer.

## INTRODUCTION

One of the leading contributors to cancer-related mortality globally is lung cancer, with predominantly non-small cell lung cancer (NSCLC) subtype (1-3). As per GLOBOCAN 2020 the latest epidemiological data, around 2.20 million new lung cancer cases were reported worldwide, attributed to 11.4% of total newly diagnosed cancer globally, accounting for 1.79 million mortality (18% of all cancer-related deaths) (4).

Lately, the finding of EGFR mutations has transformed our knowledge and treatment of lung cancer. EGFR, a tyrosine kinase transmembrane protein, plays a key role in regulating cellular proliferation and survival (5, 6). Dysregulated EGFR activation can be observed in cancer due to mutations, especially in NSCLC (35% of cases) promoting cancer growth and driving resistance to apoptosis (7). The discovery of EGFR alterations has led to targeted therapies, mainly EGFR tyrosine kinase inhibitors (TKIs), which have clinical improvement for patients with EGFR-mutants. Nevertheless, the efficacy of these treatments depends on the specific EGFR mutation type, and the onset of resistance pathway, for instance the T790M mutation is still a significant challenge (8-11). Recent reports indicate that EGFR mutations trigger unique metabolic shifts, assisting in therapeutic resistance and highly invasive cancer phenotype (12-16). Moreover, the molecular mechanisms involved in varying clinical outcomes linked with these mutations remain inconclusive.

To bridge these gaps, we conducted a structured retrospective analysis of EGFR mutational status and its correlation with clinical outcomes using a large genomic and transcriptomic dataset from tumor samples across lung, colorectal, breast, pancreatic and prostate cancers (17). Furthermore, we examined the prognostic significance among EGFR mutants in NSCLC incorporating patients survival data. Lastly, Gene Set Enrichment Analysis (GSEA) was conducted between the transcriptome of T790M and L858R variants lung cancers to understand the biological pathways that may drive therapeutic resistance and provide new therapeutic approaches.

## RESULTS

### EGFR Mutations Across Cancer Types

To classify EGFR mutations abundance across cancer types using MSK, Nature, 2024 genomic data was selected using cBioPortal (https://www.cbioportal.org) (17). Lung cancers were ranked highest with EGFR mutations incidence, followed by colorectal, breast, prostate, and pancreatic cancer with 24.2%, 2.71%, 1.29%, 0.87% and 0.71% patients respectively (Figure 1A). Lung cancer cases were further analyzed based on the mutation prevalence, the three highest EGFR mutation occurred at the site L858R in 600 patients, E746_A750del mutation in 494 cases and T790M mutation in 116 cases (Figure 1B-D).

**Figure 1.**
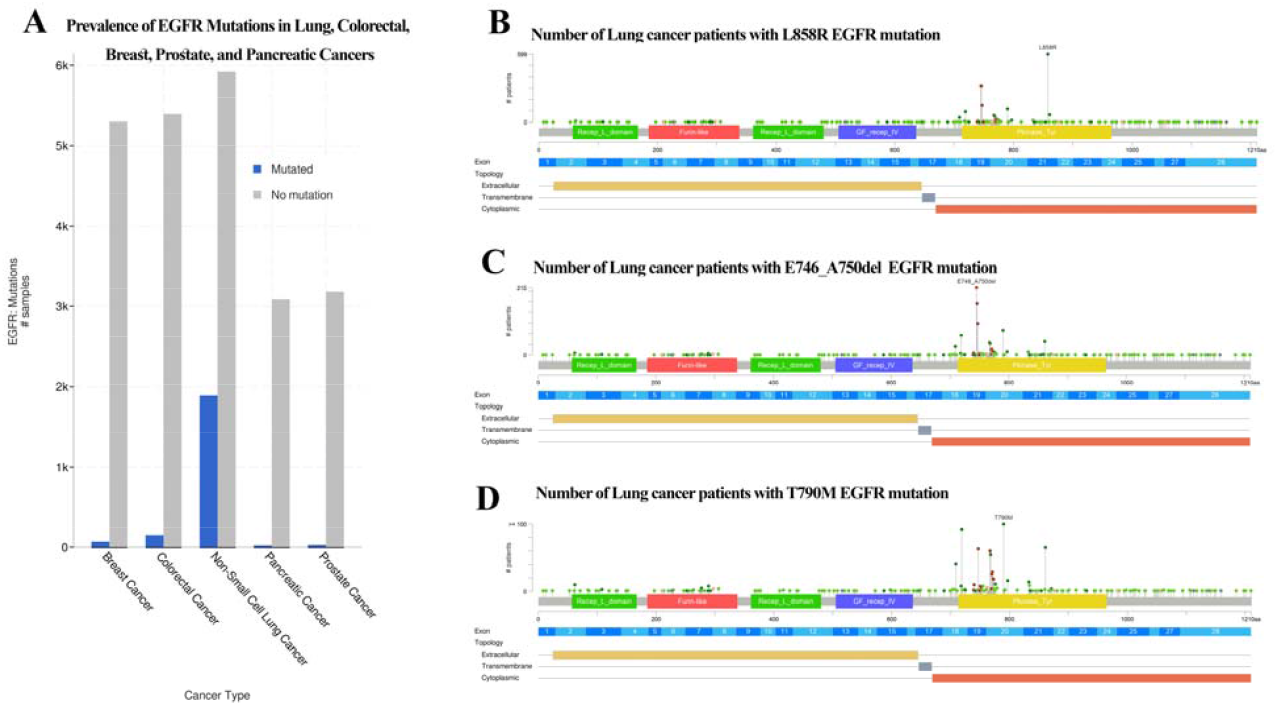
EGFR mutation frequency across major cancer types and specific mutation subtypes in lung cancer. (A) Pan-cancer analysis shows EGFR mutation rate in lungs, colorectal, breast, prostate, and pancreatic cancers. Distribution of the three most EGFR variants in lung cancer patients (B) L858R (n=600), (C) E746_A750del (n=494) and (D) T790M (n=116).

### EGFR Variants and Overall Survival

After sorting lung cancer patients based on EGFR mutation sites, overall survival rates among top three were lowest in T790M mutant patients (33.93 months median survival) (Figure 2A & 2B) and no significant difference was observed in overall survival between L858R and E746_A750del with 50.2 and 50.14 median months. The most lethal among all the EGFR mutants in Lung cancer was C797S mutants (Figure 2C) with 18.87 overall median survival months. There were other mutants which are as lethal as T790M such as L861Q, G719S, with OS of 37.22, 36.69 median months survival (Supplementary Figure 1).

**Figure 2.**
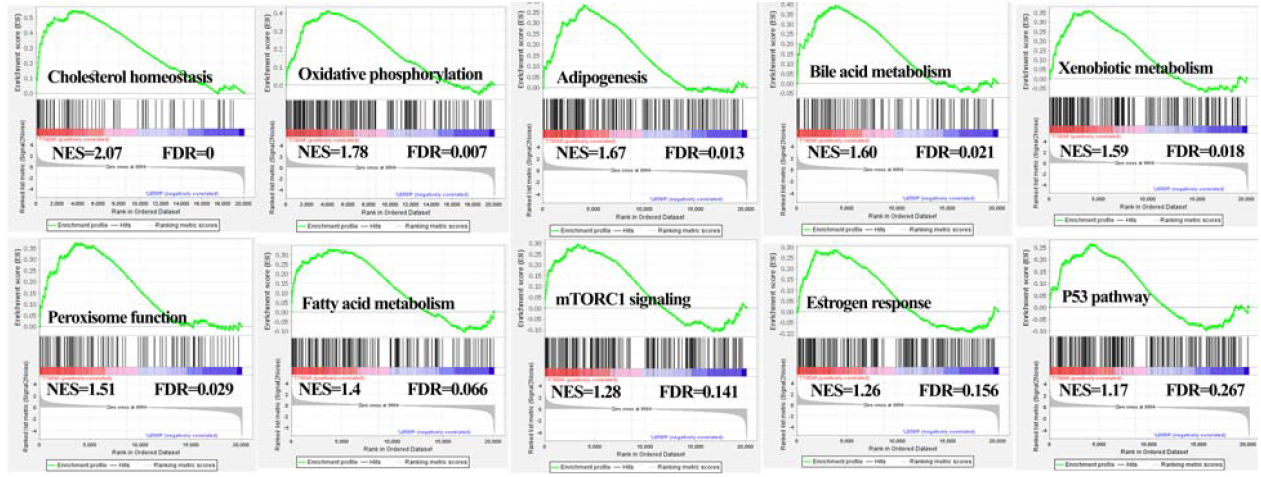
Kaplan-Meier survival analysis for lung cancer patients stratified by EGFR mutation subtypes. (A) T790M vs L858R, (B) T790M vs E746_A750del, and (C) T790M vs C797S. Statistical significance performed using log-rank test.

**Figure 3.**
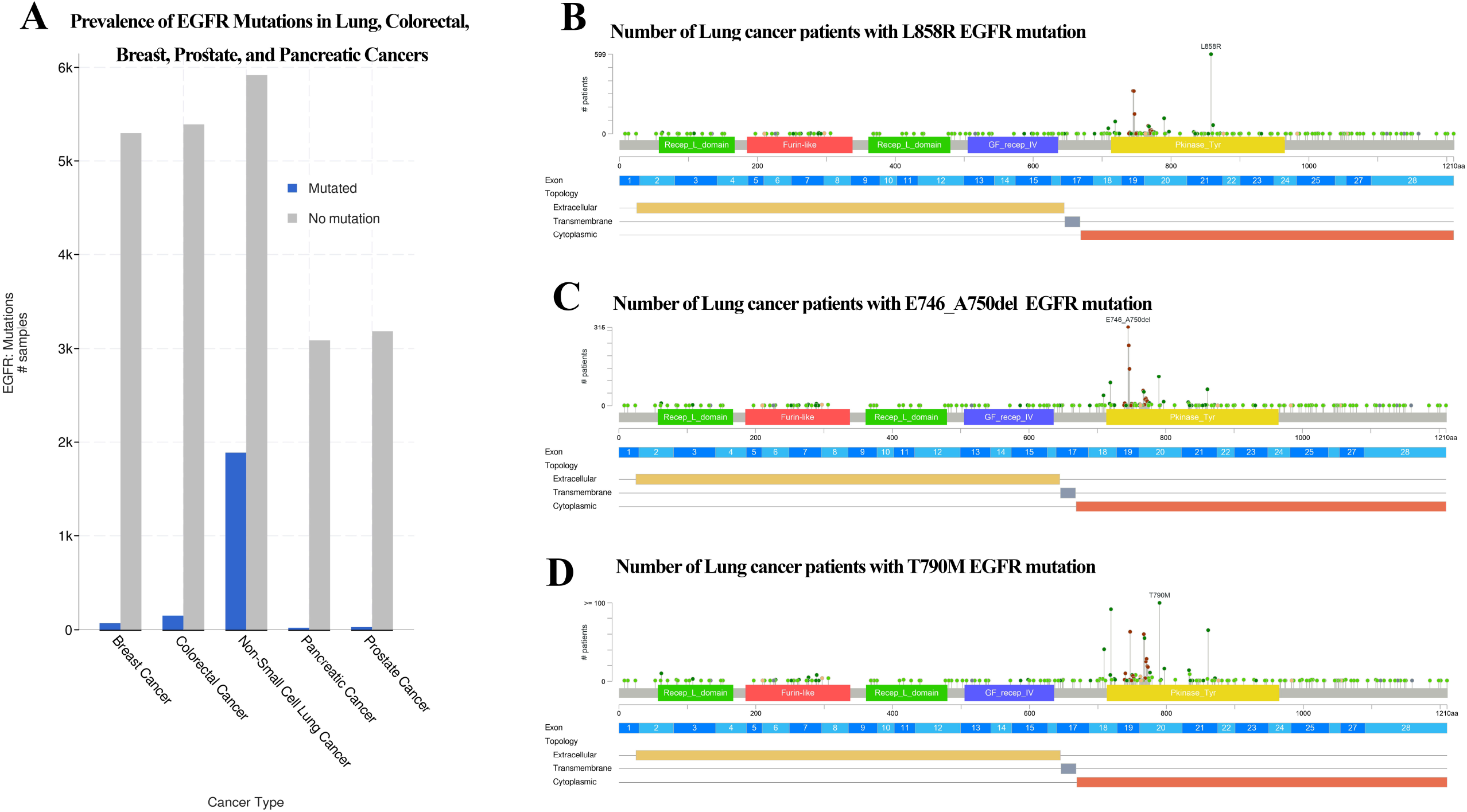
Transcriptomic data from cBioPortal analyzed by GSEA to identify enriched hallmark pathways in T790M variants.

### Metabolic Pathway Dysregulation Distinguishes T790M from L858R EGFR-Mutant Tumors

GSEA identified that T790M variants displayed vast metabolic reprogramming characterized by enrichment of a few pathways such as cholesterol homeostasis, oxidative phosphorylation, adipogenesis, bile acid metabolism, xenobiotic metabolism, peroxisome function, fatty acid metabolism, mTORC1 signaling, estrogen response (p < 0.05, NES > 1.25), and p53 pathway with modest NES of 1.18 and p value = 0.084. Assessment of key genes associated with these pathways demonstrated that T790M variants particularly overexpressed metabolic enzymes HMGCR, CPT1A, and NDUFS1, meanwhile reducing expression of regulatory genes PPARA, MTOR, SOD2, and TP53 (Supplementary Figure 2). This metabolic profile reveals that T790M mutants have adapted augmented metabolic flexibility and detoxification potential because of enhanced oxidative activity and lipid metabolism, along with diminished antioxidant defense and regulatory control, possibly elucidating their link with treatment resistance and poor prognosis compared to L858R variant cases.

## DISCUSSION

EGFR mutations are genetic changes that can result in aberrant activation of EGFR signaling pathways, driving neoplastic growth and cancer development (18, 19). EGFR mutations play a major role in the pathogenesis of NSCLC, predominantly in the adenocarcinoma subtype. These mutations, frequently identified as exon 19 deletions and exon 21 L858R point mutations, lead to constitutive activation of the EGFR tyrosine kinase domain (20-22).

This retrospective study examines EGFR mutational landscapes across various cancer types, revealing lung cancer as the predominant malignancy containing EGFR mutations which is consistent with previous epidemiological and molecular studies (23-26), with L858R and E746_A750del representing most frequent variants. Our findings displayed significant prognostic heterogeneity within EGFR variants, with C797S cases conferring the worst survival rate (median OS: 18.87 months), whereas T790M mutations revealed second worst clinical outcome (median OS: 33.93 months). While L858R or E746_A750del mutations achieved significantly longer median overall survival (∼50 months), signifying these mutants may represent more therapeutically targetable phenotypes. These discoveries align with standard disease patterns, where T790M is known as a resistance mutation to 1^st^ and 2^nd^ generation EGFR tyrosine kinase inhibitors (TKIs), and C797S acquire resistance to 3rd-generation drugs like Osimertinib (13, 27-30). These disparate survivals highlight the need for precision medicine approaches personalized to specific EGFR mutants, with C797S-positive cases demonstrating a mostly high-risk population possibly requiring novel therapeutic modalities, including emerging 4^th^ generation TKIs or therapeutic combinations under clinical studies.

One of the major highlights our analysis is the distinct metabolic reprogramming linked with the T790M mutation. GSEA revealed that T790M cases are characterized by upregulation of pathways involved in cholesterol homeostasis, oxidative phosphorylation, fatty acid metabolism, and detoxification processes. This metabolic pattern is corroborated by upregulation of important metabolic enzymes such as HMGCR, CPT1A, and NDUFS1, which contribute to Bioenergetics and lipid biosynthesis. Meanwhile, the reduced expression of regulatory and tumor suppressor genes such as PPARA, MTOR, SOD2, and TP53 indicate a disruption of metabolic and genomic regulation, possibly causing these variants more tolerance to therapeutic stress with increased aggressiveness and resistance to standard EGFR-targeted therapies.

The enriched p53 pathway in T790M patients, although low levels of TP53, suggests a complex interplay of regulatory network where activation of downstream pathway may occur independently of p53. This could reflect compensatory stress responses or alternative regulatory pathways that attempt to preserve cellular homeostasis in the context of oncogenic stress yet fail to provide effective tumor suppression due to the lack of functional p53. Our results are consistent with emerging evidence that metabolic flexibility represents a hallmark of drug-resistant cancer cells and could be a key factor in their aggressiveness (13, 31, 32). However, the retrospective study design and dependence on bulk RNA sequencing data needs a cautious interpretation. The biological activity of these metabolic pathways in T790M requires additional experimental support using single cell RNA sequencing.

Overall, our analysis highlights the intratumorally heterogeneity of EGFR-mutant in lung cancer and the importance of integrating mutation-specific survival data and pathway signatures into clinical decision-making to optimize patient stratification and enable precision targeting of resistance mechanisms. To summarize, the metabolic and molecular profiling of EGFR mutants, particularly T790M, offers a basis for developing next-generation targeted therapies and emphasizes the ongoing challenges in managing advanced NSCLC.

## METHODS

### Evaluation of EGFR Mutational Status Across Multiple Cancer Types

We analyzed the publicly available oncologic real-world dataset (MSK, Nature, 2024) by cBioPortal (https://www.cbioportal.org) to access EGFR gene mutations in diverse cancers. The dataset comprised 25,040 samples representing breast, prostate, non-small cell lung cancer (NSCLC), pancreatic, and colorectal cancers. We evaluated the mutational status of EGFR in these samples and identified the most frequently altered EGFR mutation site across the different cancer types. The distribution of the highest-frequency EGFR site mutations was visualized for comparative analysis.

### Overall Survival Investigation of EGFR-Mutant Lung Cancer

To study the mutational status of the EGFR and its effect on OS in lung cancer patients, EGFR mutation data were filtered to download only lung cancer from cbioportal. Survival data were procured from the clinical outcome annotations presented in the dataset. Kaplan–Meier survival analysis was performed using GraphPad application to compare OS between patients with different EGFR mutation subtypes. Statistical significance was determined using the log-rank test, and p-values < 0.05 were considered statistically significant. The raw data analyzed for this study is provided in Supplementary File 1.

### GSEA Using Hallmark Pathways in EGFR Mutant Subgroups of Lung Cancer

GSEA was performed to compare clinical transcriptomic profiles between EGFR T790M and L858R mutant in lung cancer (due to lack of transcriptome data for C797S, L861Q, and G719S we could not perform the GSEA for those variants). The analysis was conducted using the Molecular Signatures Database (MSigDB) gene set collection h.all.v2024.1.Hs.symbols.gmt, which covers the Hallmark gene sets. The analysis was conducted with a total of 1,000 permutations, with the permutation type set to “gene set”. The rest of the parameters were kept at their default settings. The raw data analyzed for this study is provided in Supplementary File 2.

## AUTHOR CONTRIBUTIONS STATEMENT

TK and MT planned and designed the study; TK processed the raw data (transcriptome and genome); TK performed GSEA; TK and MT analyzed the data; TK wrote the original draft; TK and MT reviewed and edited the final manuscript and approved submission.

