## Supplementary figures and images for "EGFR Mutation Subtypes Modulate Distinct Metabolic Profiles and Clinical Outcomes in Lung Cancer: A Retrospective Analysis"

### Supplementary Figure 1

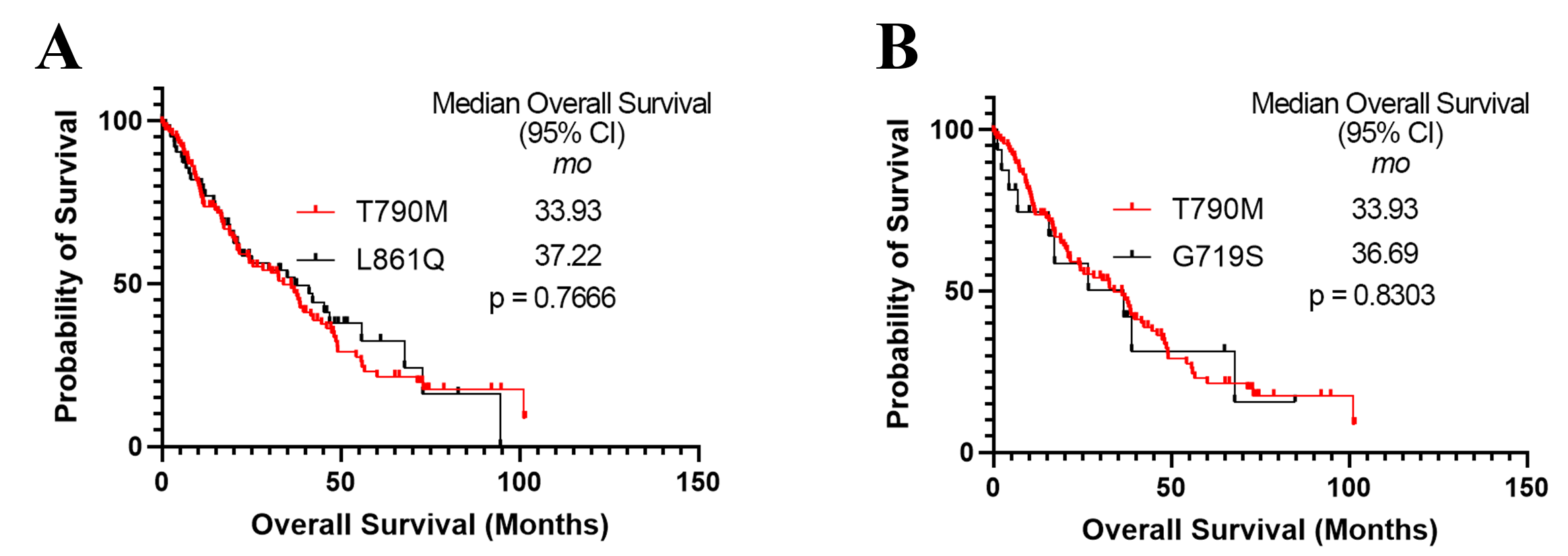

### Supplementary Figure 2

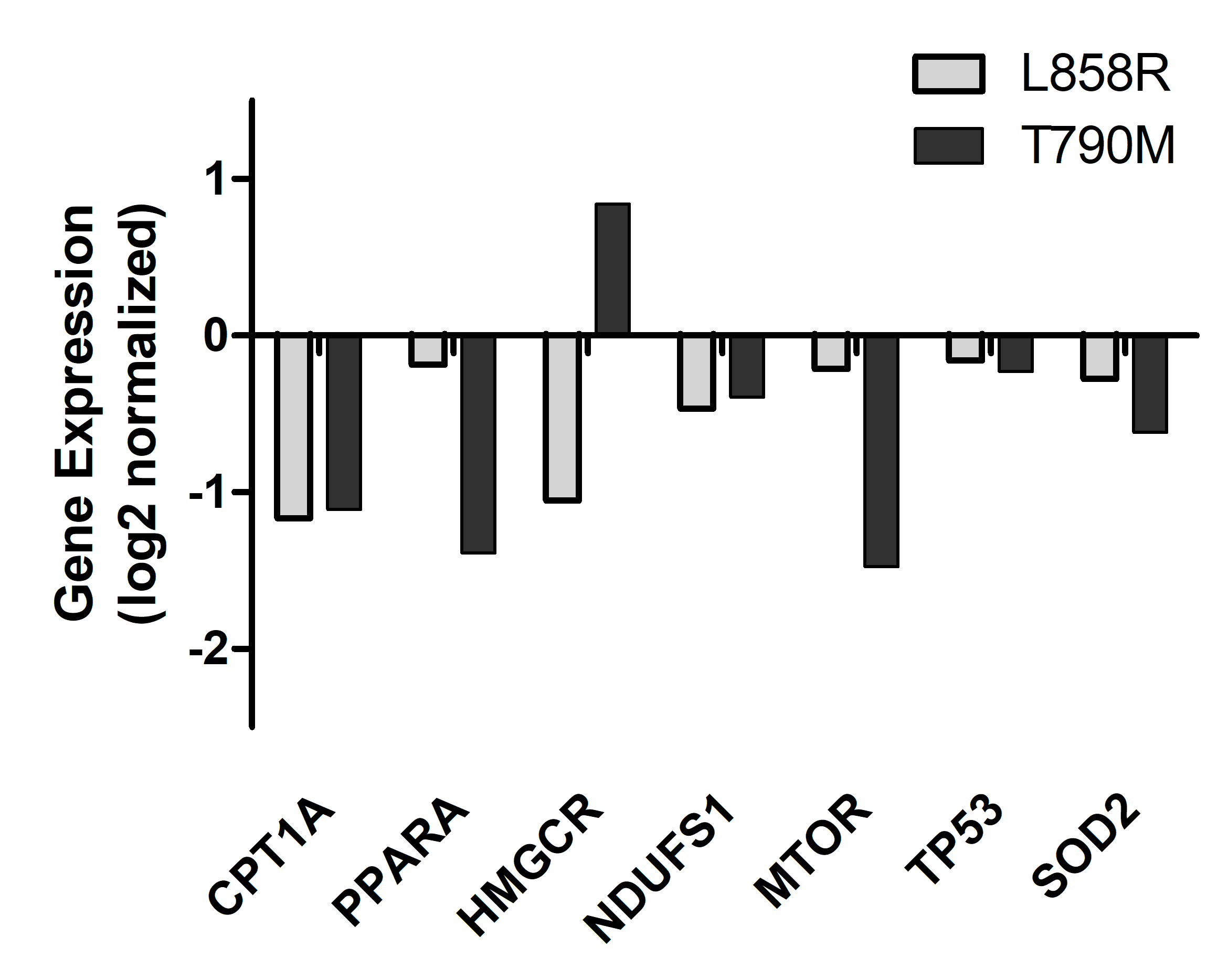
